# Improving AlphaFold3 by Engineering MSA and Template Inputs

**DOI:** 10.64898/2026.04.22.720119

**Authors:** Pawan Neupane, Jian Liu, Jianlin Cheng

## Abstract

AlphaFold3 introduces a unified framework for predicting the structures and interaction of several biomolecules including single-chain protein monomers, multi-chain protein multimers, and protein–ligand complexes. While it achieves the state-of-the-art performance in most predictions, its prediction accuracy depends on the quality of multiple sequence alignment (MSA) and structural template inputs. There are few works of using customized MSAs and templates to improve AlphaFold3. In this work, we systematically investigate how diverse and carefully engineered MSAs and templates can be leveraged to improve AlphaFold3 predictions. We evaluate our methods on protein monomers, multimers, and protein-ligand complexes, and observe consistent, sizable gains in structure prediction accuracy for monomers (TM-score 0.937 vs 0.882), for multimers (DockQ score 0.550 vs 0.525), and for protein-ligand complexes (ligand RMSD 3.258 Å vs 4 Å) compared to the default AlphaFold3. Moreover, for the first time, we demonstrate that AlphaFold3 performs significantly better than AlphaFold2 when both use the same customized MSA and template inputs. The results highlight the importance and effectiveness of using diverse MSAs and templates to improve AlphaFold3.

## 1 INTRODUCTION

Deep learning has fundamentally reshaped structural biology by enabling accurate prediction of three-dimensional biomolecular structures directly from sequence information since it was introduced in the field in 2012 [1]. A major milestone was the development of transformer-based AlphaFold2 [2], which achieved near-experimental accuracy for predicting the tertiary structures of most single-chain proteins (monomers) and established deep learning as the dominant paradigm for computational structural biology. Subsequent developments extended these approaches to more complex modeling scenarios, including modeling protein complexes and interactions [3, 4, 5, 6] and leveraging the evolutionary information learnt by protein language models [7]. Most recently, AlphaFold3 [8] and RoseTTAFoldAll-Atom [9] introduced unified deep learning frameworks for modeling proteins, nucleic acids, ligands, and their interactions, expanding the scope of deep learning prediction to the structures of almost all the major biomolecules and their interactions. Particularly, AlphaFold3-based predictors achieved the state-of-the-art performance in most prediction categories during the 16th Critical Assessment of Techniques for Protein Structure Prediction (CASP16) in 2024 [10, 11, 12, 13, 14, 15, 16].

The accuracy of deep learning-based protein structure prediction depends on input quality, relying primarily on evolutionary information in multiple sequence alignments (MSAs) and, to a much lesser degree, on structural templates.[5, 6, 17, 18, 16, 14]. MSAs capture patterns of residue conservation and coevolution arising from structural and functional constraints, providing a critical link between sequence variation and three dimensional structure [19, 20]. In AlphaFold-based prediction, such signals influence both local structural features and long-range residue relationships. As a result, prediction accuracy can be sensitive to the depth, diversity, and composition of the input MSAs, as well as to the availability and quality of structural templates, particularly for difficult targets.

However, because no single MSA construction method is guaranteed to generate better MSAs for all protein targets and it is still impossible to accurately estimate the quality of MSAs for structure prediction, recent research sought to use multiple MSAs generated by different alignment methods to improve protein structure prediction [21, 17, 22, 14]. Particularly, some top predictors in the CASP15 and CASP16 experiments such as MULTICOM4 [14] demonstrated that using diverse MSAs generated by complementary MSA construction methods can substantially improve the accuracy of AlphaFold2-based protein tertiary and quaternary structure prediction [17, 22, 14]. In addition, diversification of template input can also have some positive impact on structure prediction for some targets [21, 17, 22, 14]. And using diverse MSAs and templates is more effective in improving AlphaFold2 prediction than simply increasing the number of structural models predicted from a single input [14].

However, despite the substantial evidence indicating that MSA and template diversification strategies can enrich evolutionary and structural information to substantially improve the prediction accuracy of AlphaFold2, their roles have not been systematically examined in the context of AlphaFold3. To the best of our knowledge, there is only one work [23] showing that using an unpaired strategy to generate MSAs for protein complexes without pairing the MSAs of individual chains can help AlphaFold3 produce more accurate structure prediction for some protein complexes than its default paired MSA strategy. But it is still unclear if diverse MSA and template inputs generated by different alignment techniques and from different protein databases can substantially improve the performance of AlphaFold3 as it does for AlphaFold2, how sensitive AlphaFold3 predictions are to alternative choices of MSAs (e.g., sequence alignment-based MSAs or structure alignment-based MSAs), how these effects compare to those observed in AlphaFold2 under the same matched conditions, and whether the benefits of MSA and template diversification generalize across distinct structural regimes, including protein monomers, protein multimers, and protein–ligand complexes.

Moreover, even though the CASP16 prediction results of some top predictors such as our MULTICOM4 system built on top of AlphaFold2 and 3 showed that AlphaFold3 using only the default MSA and template performed slightly better than or similarly to AlphaFold2 using multiple diverse MSA and template inputs, it is still unknown how much advantage that AlphaFold3 may have over AlphaFold2 if both of them using the same MSA and template diversification strategy.

To answer these questions, in this work, we implement the same MSA and template diversification strategy that improves AlphaFold2 in our CASP16 MULTICOM4 protein structure prediction system [16, 14] to generate the inputs for AlphaFold3. We blindly applied AlphaFold3 equipped with the MSA and template enhancement strategy (called Custom AF3) to CASP16 targets and directly compared its prediction results with those of AlphaFold2 with the exactly same strategy (called Custom AF2) obtained during the CASP16 experiment.

Specifically, Custom AF3 uses dozens of complementary sequence alignment-based MSAs generated by multiple sequence search tools and expanded protein sequence databases as well as structure-alignment based MSAs generated by FoldSeek[24]-based alignment of predicted structures to the structures in Protein Data Bank[25] and AlphaFoldDB [26] as input. Each MSA together with the default structural templates or templates found by the customized template search tools is used as input for AlphaFold3 to generate a number of structural models. The structural models for all the different inputs are pooled together as final predictions of Custom AF3.

We assessed the prediction accuracy of Custom AF3 across three classes of biomolecules: protein monomers, protein complexes (multimers), and protein–ligand complexes, which were used as targets in the CASP16 experiment. Across hundreds of CASP16 monomer, multimer, and protein-ligand complex targets, we observe a consistent and significant performance improvement of using diverse MSAs and templates with AlphaFold3 over the default input.

Moreover, our experiment shows that, with the same, diverse MSAs and templates as input, AlpahFold3 performs significantly better than AlphaFold2. It is worth noting that the customized MSAs and templates for CASP16 targets used by Custom AF3 and AF2 in our benchmark were generated during the 2024 blind CASP16 experiment before the true structures of the targets were known. Therefore, the comparison between the two is fair and objective without any data leakage. Together, these findings firmly establish MSA and template diversification as a broadly effective strategy for improving AlphaFold3 predictions for protein monomers, multimers, and protein-ligand complexes and provide a simply, yet highly effective approach to enhancing AlphaFold3.

## 2 MATERIALS AND METHODS

### 2.1 CASP16 Datasets

We evaluated AF3 with the enhanced MSA and template generation (Custom AF3), default AF3, and AF2 with the enhanced MSA and template generation (Custom AF2) on the datasets of the 2024 CASP16 experiment. The protein monomer dataset includes 12 targets that exist exclusively as monomers (i.e., not as subunits of protein complexes). The protein multimer dataset includes 36 targets, excluding H1217, H1227, and H1272 due to their large molecular sizes that exceed the available GPU memory limits. The protein–ligand complex dataset includes four superligand targets (L1000, L2000, L3000, and L4000) for which experimentally determined native structures are available. The protein in each superligand target interact with many ligands, including small-molecule ligands, peptide ligands, and metal or cofactor ions. Each distinct protein–ligand pair was treated as an independent target, resulting in a total of 286 protein–ligand complex targets for which appropriate native structures were made available following the competition.

### 2.2 Experimental Design

To evaluate the impact of multiple sequence alignment (MSA) construction and template selection on prediction performance, we established a controlled benchmark for AF3 and AF2 on the CASP16 datasets described above. In this benchmark, MSA and template inputs were systematically varied, while all other inference parameters (e.g., number of recycles and model settings) were kept fixed to isolate the effects of input variation.

For each CASP16 target, multiple MSAs and structural templates generated by MULTICOM4 during the 2024 CASP16 experiment were used to construct distinct MSA–template combinations. Because these inputs were produced before the release of experimental structures, the evaluation strictly reflects blind prediction conditions and avoids any potential data leakage. Each MSA–template combination was provided as input to AF3 and/or AF2 to generate structural predictions, allowing a direct comparison of the effects of alternative alignment and template choices on them.

For AF3, predictions were generated using the same model version (v1) with default inference parameters across all runs, ensuring that observed performance differences could be attributed solely to variations in MSA and template inputs. For AF2, predictions were generated using the standard model weights (`monomer` or `multimer_v3`), with identical MSA and template inputs supplied to ensure direct comparability between AF2 and AF3. Depending on target size and GPU resource availability, between 25 and 100 models were generated per MSA–template combination for each target.

### 2.3 Implementation of AF3 with enhanced MSA and template generation (Custom AF3)

#### 2.3.1 Engineering MSA and template input for protein monomer

Fig. 1 illustrates the workflow of Custom AF3. To capture a broad range of evolutionary constraints, the MSA and template engineering module [16] in Custom AF3 constructs a diverse set of multiple sequence alignments (MSAs) for a protein monomer and uses different tools and templates to select structural templates. It combines multiple complementary MSAs from different search strategies and databases with different template options to form a diverse set of MSA-template combinations as inputs for AF3.

**FIGURE 1.**
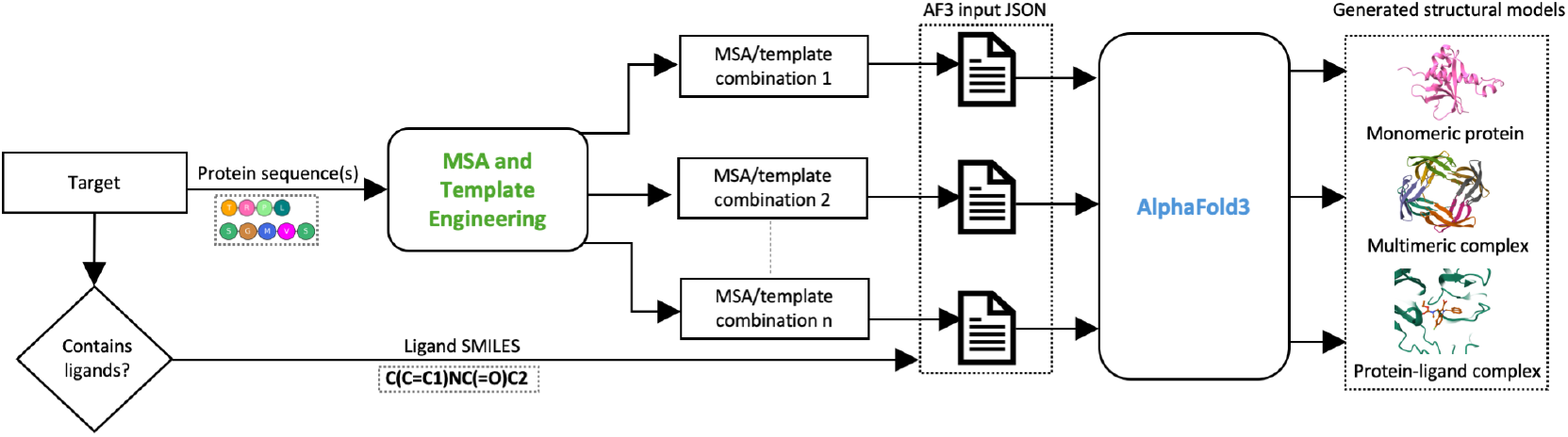
The structure prediction pipeline of Custom AF3. For a target (protein monomer, protein complex, or protein-ligand complex), protein sequence(s) are processed by the customized MSA and template engineering module in MULTICOM4 [18, 16] to generate diverse MSA and template combinations, which are encoded as AlphaFold3-compatible JSON inputs. For protein–ligand complex, ligand SMILES strings are directly included in the JSON alongside receptor protein inputs. The input of the target is used by AlphaFold3 to generate a user-specified number of structural models.

It first generates default MSAs following the paradigm used in the default AF3, where the target sequence is searched against both general-purpose protein databases (UniRef30 [27], UniRef90 [28]) and metagenomic databases (BFD [29, 30, 31], MGnify [32]) using HHblits [33] and JackHMMER [34]. It is worth noting that the default AF2 MSAs used in this work were generated in the similar way during the CASP16 experiment [16].

It then generates additional MSAs using DeepMSA2 [22] and Dense Homolog Retriever (DHR) [35]. DeepMSA2 iteratively searches a combination of general and metagenomic resources, including Metaclust [30], TaraDB [36], JG-Iclust [37], and MetaSourceDB [38], using both HHblits and JackHMMER, and produces multiple complementary alignment variants (e.g., qMSA, dMSA, hMSA). It also retains intermediate MSAs generated during the iterative search process, resulting in a total of 15 DeepMSA2-related MSA variants. In contrast, DHR leverages protein language model embeddings to retrieve remote homologs, which is beneficial for targets with limited close relatives and can increase effective alignment depth. Furthermore, Custom AF3 also includes MSAs generated by ColabFold [21].

Beyond retrieved homologs in the protein sequence databases, it uses a fine-tuned ESM-2 [7] to generate synthetic homologs to enhance MSAs. Specifically, residues in a target sequence are systematically masked and replaced with high-confidence predictions made by the fined-tuned ESM-2, and the resulting pseudo-homologous sequences are appended to the default MSA to form an extended alignment (“ESM-MSA”). The extended MSAs can improve robustness when naturally occurring homologs are sparse.

For template identification, it uses the MSA generated by searching UniRef90 with JackHMMER as input for HHsearch to build a profile. HHsearch searches the profile against two template libraries: pdb70_v2024_03 (the standard AlphaFold2 template database clustered at 70% sequence identity) and PDB_sort90 [18] (an in-house curated database clustered at 90% sequence identity) to retain broader structural diversity for similar structural templates.

Because using templates does not always help structure prediction, it also includes a no-template setting in which template features are not provided (set to null).

Overall, combining the MSAs with alternative template options above yields up to 22 distinct MSA-template combinations for protein monomers (see a summary of some representative monomer MSA-template combinations in Supplementary Table S1).

#### 2.3.2 Engineering MSA and Template Input for Protein Multimer

For a protein multimer, Custom AF3 constructs both sequence alignment-based and structure alignment-based MSAs for each unique subunit (chain) of the multimer first. Sequence alignment-based MSAs are generated using the same procedures described in Section 2.3.1. Structure-based MSAs are obtained by using FoldSeek [24] to search a top-ranked tertiary structure model of the subunit selected by global pLDDT against pdb_complex [17], AlphaFoldDB [26] and the ESM Metagenomic Atlas [7]. The resulting structural alignments between the structural model and the similar hits are converted into structure alignment-based MSAs, providing complementary MSAs beyond sequence-only searches because structure alignments can find remote homologs that cannot be recognized by sequence alignment.

To capture inter-chain residue-residue co-evolution, it constructs multimer-level MSAs by pairing and concatenating subunit MSAs. For sequence-based multimer MSAs, subunit alignments are concatenated using multiple pairing signals, including species information [3, 39], UniProt accession IDs, STRING interactions [40], and known PDB complexes. This yields multiple concatenated MSA variants (13 for heteromers and 7 for homomers, with homomers excluding STRING- and accession-based pairing). It additionally applies the DeepMSA2 pairing protocol, in which DeepMSA2-generated subunit MSAs are ranked by the best AlphaFold2 pLDDT score achieved using each MSA to generate structural models and the top-ranked MSAs are concatenated, producing up to 20 multimer MSA variants depending on subunit availability. For structure alignment-based multimer MSAs, structure-derived alignments that share the same identifier are concatenated across subunits, while unpaired alignments are retained to preserve complementary structural signals.

Because pairing monomer MSAs is not always beneficial (e.g., for antibody targets), the unpairing strategy of simply concatenating monomer MSAs without pairing them is also used to generate some multimer MSAs.

Templates are identified at the subunit level using HHsearch [41] search against three template libraries: PDB70, pdb_complex [17], and pdb_sort90 [18]. Structural templates are also identified by using each subunit’s predicted structure as input to FoldSeek to search against pdb_complex. Multimer templates are then assembled by concatenating subunit templates that share the same PDB code; if fewer than four concatenated templates are available, top-ranked unpaired subunit templates are added to ensure at least four templates per multimer. In addition, pdb_seqres from the PDB (the default template database for AF2 and AF3) is used to retrieve templates without multimer-template concatenation. Similar to protein monomers, a no-template setting (setting template to null) is also used for protein multimers.

Overall, the constructed multimer MSAs and assembled multimer templates produce up to 62 distinct MSA-template combinations for AF3 to generate structural models. (see a summary of some representative multimer MSA-template combinations in Supplementary Tables S2 and S3).

#### 2.3.3 Engineering MSA and Template Input for Protein-Ligand Complex

For a protein-ligand complex target, the receptor protein is processed using the same MSA engineering pipelines described above for monomeric or multimeric proteins, depending on whether the receptor is a monomer or multimer.

Ligands are represented as the canonical SMILES strings, which are appended to the MSA and template input of the receptor to form the complete protein-ligand complex input compatible with AF3. No ligand-specific MSA is constructed. Instead, the influence of the ligand on structure prediction is modeled directly through its chemical representation contained in the input. This simple, unified treatment allows protein–ligand prediction to benefit from the same enriched protein MSA and template information used for protein-only targets, enabling AF3 to jointly model protein sequence, structure, and ligand chemistry.

### 2.4 Evaluation Metrics

For assessing the accuracy of predicted protein monomer structures, we employed four widely used metrics: TM-score and Global Distance Test - Total Score (GDT-TS) for global fold accuracy, Global Distance Calculation for Side-chains (GDC-SC) for side chain accuracy, and Local Distance Difference Test (lDDT) for computing local distance differences of all atoms (Fig. 2a). TM-score was computed using the TM-score program [42, 43], which provides a length-independent measure of structural similarity between a predicted model and its corresponding experimental structure. GDT-TS and GDC-SC scores were calculated using the `lga` tool (Local–Global Alignment) [44], following the standard CASP evaluation protocol, whereas lDDT was calculated using the implementation [45] in the `OpenStructure` framework [46].

**FIGURE 2.**
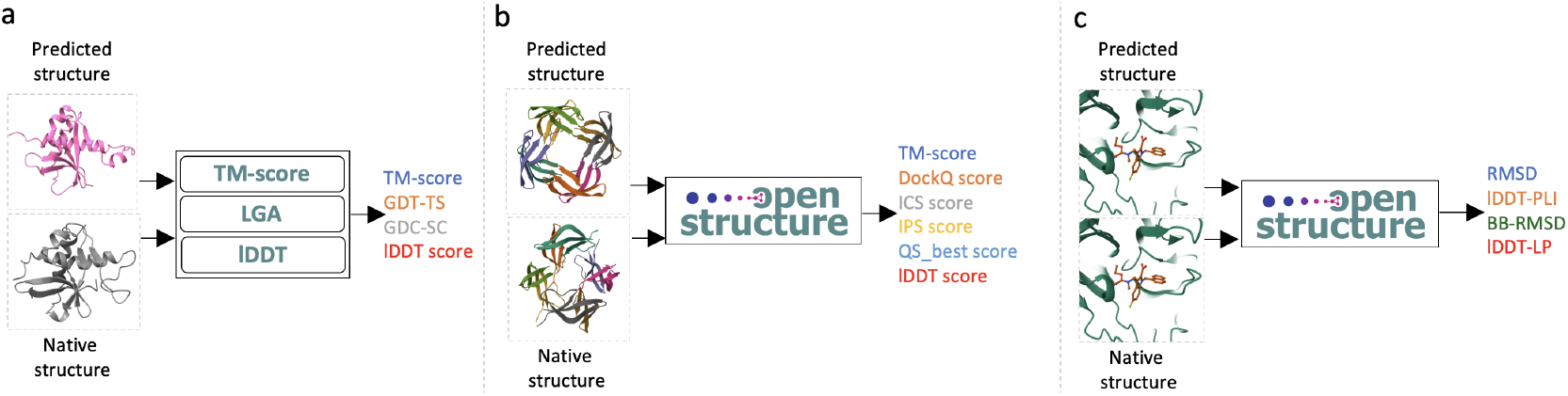
Structure evaluation pipelines for three types of targets. **a**. Protein monomer evaluation using `TM-score` [42, 43], `LGA` [44], and `lDDT` [45] programs reporting global (TM-score and GDT-TS), side-chain (GDC-SC) and local-level (lDDT) quality metrics. Three tools (TM-score, LGA, and lDDT) are used to compare a predicted structure with the experimental structure to obtain these scores. **b**. Protein multimer (complex) evaluation using `OpenStructure`[46], reporting global (TM-score), interface-level (DockQ score, ICS score, IPS score, QS_best score) and local (lDDT score) metrics. **c**. Protein–ligand evaluation using `OpenStructure` with ligand-position-specific (RMSD) and protein-ligand-interface-specific (lDDT-PLI),backbone-confirmation-specific (BB-RMSD) and local-geometry-specific (lDDT-LP) quality metrics.

To evaluate the accuracy of predicted protein multimer structures, we employed the `compare-structures` pipeline of the `OpenStructure` framework [46], which provides a unified framework for structural comparison and interface quality assessment (Fig. 2b). Using this framework, we computed a comprehensive set of metrics: TM-score for global structural similarity, DockQ for assessing docking quality, ICS (Interface Contact Similarity), IPS (Interface Patch Similarity) for capturing interface patch correspondence, QS_best for quaternary structure similarity, and lDDT for local distance agreement independent of superposition. Together, these complementary measures provide a balanced evaluation of both global fold correctness and interface-specific accuracy, enabling a thorough assessment of multimeric complex predictions.

For protein–ligand complexes, we applied the `compare-ligand-structures` pipeline available in the `OpenStructure` framework [46] to assess both global and local accuracy (Fig. 2c). This pipeline computes multiple complementary metrics: ligand RMSD to measure the overall positional accuracy of the ligand with respect to the experimental structure, lDDT_PLI to quantify local distance deviations at the protein–ligand interface, backbone RMSD (BB-RMSD) to assess conformational changes in the protein backbone upon ligand binding, and lDDT_LP to evaluate local geometry around the ligand. Together, these metrics capture binding pose fidelity, local interface quality, and structural perturbations of the receptor, enabling a comprehensive evaluation of protein–ligand complex predictions.

## 3 RESULTS AND DISCUSSION

Predictions were evaluated on the valid CASP16 targets (2.1) under the controlled experimental settings (2.2), including protein monomers, protein multimers, and protein–ligand complexes. AF3 was applied to all three target types, whereas AF2 was evaluated on protein monomers and multimers only, as it does not support ligand prediction. For each comparison, the same number of models was generated per target and per input condition to ensure fairness and consistency in performance assessment. Particularly, we focused on the following two comparisons: (1) AF3 with multiple MSA-template combinations (Custom AF3) versus AF3 with default MSA and template input (default AF3), and (2) AF3 versus AF2 with the same customized inputs (Custom AF3 versus Custom AF2).

### 3.1 Tertiary Structure Prediction for Protein Monomers

#### 3.1.1 Comparison of Custom AF3, default AF3, and Custom AF2 on CASP16 monomer targets

We compared Custom AF3, default AF3 and Custom AF2 on 12 standalone CASP16 monomer targets that are not a part of any protein complex (Fig. 3), excluding monomer targets that are units of a protein multimer because their tertiary structure prediction accuracy is mostly determined by the quality of multimer structure prediction. The default AF3 used the standard MSA and template input generated by AlphaFold3 to predict 400 to 2100 structural models per target (equal to 25–100 models multiplied by the number of MSA–template combinations). In contrast, the Custom AF3 generated the same total number of structural models, but used all custom MSA–template combinations, producing 25–100 models for each combination. Both of them use the AF3 ranking score to rank and select models for evaluation. Similar to the Custom AF3, the Custom AF2 also used all Custom MSA–template combinations to generate a large number of structural models; however, to ensure a fair comparison, an equal number of models (25–100) from each combination were randomly sampled and ranked using the global pLDDT score.

**FIGURE 3.**
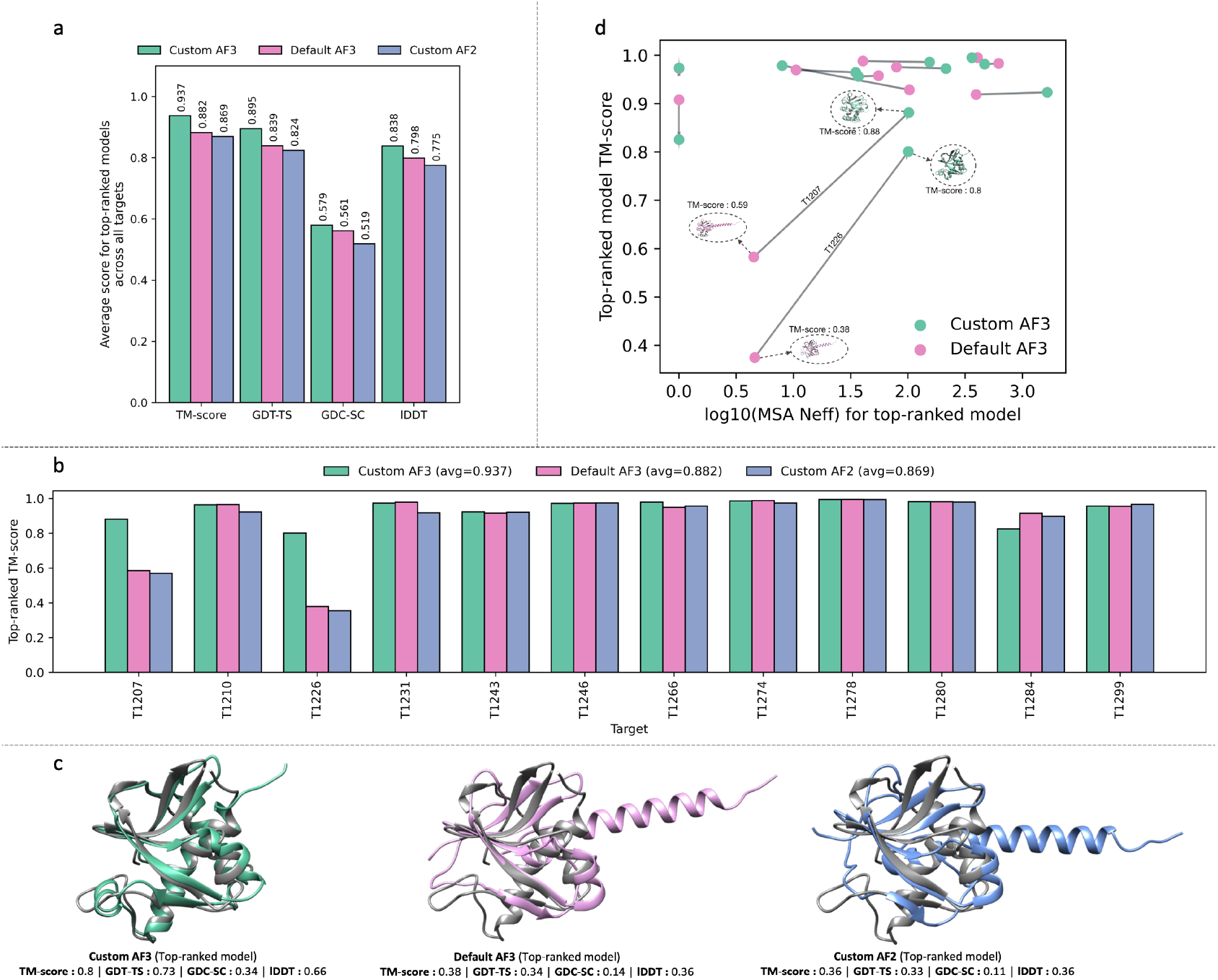
Comparative performance of Custom AF3, default AF3, and Custom AF2 on CASP16 monomer targets. **a**. Average per-target performance in terms of TM-score, GDT-TS, GDC-SC, and lDDT for the top-ranked model from each method. **b**. TM-scores of top-ranked models of Custom AF3 (green), default AF3 (pink) and top-ranked Custom AF2 (blue) for 12 monomer targets. Mean TM-scores across all targets are shown in parentheses. **c**. The top-ranked models for target T1226 generated by each method superimposed with the native structure. The native structure is shown in silver, and predicted models are colored by method, illustrating differences in structural accuracy. **d**. Relationship between MSA depth, measured by the effective number of sequences (Neff, scaled logarithmically), and TM-score of top-ranked models of Custom AF3 and default AF3 across all targets. Two dots (green and pink) denoting the Custom AF3 model and the default AF3 model for the same target are connected for direct comparison.

Prediction quality was evaluated by four complementary metrics that assess different aspects of structural accuracy, including global structural quality scores (TM-score [42, 43] and GDT-TS [44]), side-chain accuracy measured by GDC-SC (Global Distance Calculation for side chains), and local structural quality scores (lDDT [45]).

The average scores for top-ranked models across the 12 monomer targets show that Custom AF3 (TM-score : 0.937, GDT-TS : 0.895, GDC-SC : 0.579, and lDDT : 0.838) consistently outperforms default AF3 (TM-score : 0.882, GDT-TS : 0.839, GDC-SC : 0.561, and lDDT : 0.798) and Custom AF2 (TM-score : 0.869, GDT-TS : 0.824, GDC-SC : 0.519, and lDDT : 0.775) (Fig. 3a). It interesting to observe that default AF3 using default MSAs and templates performs better in tertiary structure prediction than Custom AF2 with diverse MSAs and templates, and Custom AF3 with diverse MSAs and templates further expands the advantage over Custom AF2. And the improvement in terms of some metrics such as TM-score is pronounced (0.937 of Custom AF3 vs 0.882 of default AF3 vs 0.869 of Custom AF2). We also evaluated the three methods using the quality score of the best structural models produced by each method. The results are provided in Supplementary Table S4, which show a similar trend.

Across individual targets, the top-ranked models from Custom AF3 achieve higher TM-scores than the other two methods (Fig. 3b) for many of them, with improvements particularly substantial on two hard targets (T1207 and T1226). For T1226, default AF3 and Custom AF2 failed to predict the correct fold (TM-score of top-ranked model < 0.5) and for T1207 they barely generated a top-ranked model with correct fold (0.5 < TM-score of top-ranked model < 0.6). In contrast, for both cases, Custom AF3 generated a high-quality top-ranked model with TM-score >= 0.8. These results show that it is particularly useful to use diverse MSAs and templates with AF3 for hard monomer targets.

Fig. 3c visualizes the difference in the top-ranked structural models predicted by Custom AF3, default AF3 and Custom AF2 for Target T1226 in comparison with its native (true) structure. Only the top-ranked Custom AF3 model has a correct overall fold (TM-score: 0.8), while the other two top-ranked models, default AF3 (TM-score: 0.38) and Custom AF2 (TM-score: 0.36) only correctly folded the N-terminal region but incorrectly folded the C-terminal region as an extended helix.

We examined how the MSA diversification strategy of Custom AF3 may change MSA depth and model accuracy in comparison with default AF3. Fig. 3d plots the TM-score of the top-ranked model against the effective number of sequences (Neff, scaled logarithmically to incorporate all the targets in the figure) in the MSA used to generate the top-ranked model for each target for Custom AF3 and default AF3 respectively. In two hard cases (T1207 and T1226), the Custom MSA has a much higher depth than the default MSA, which helps move the prediction accuracy from low or medium TM-scores to the high scores (TM-score ≥ 0.8). The results demonstrate that using multiple MSAs has a higher chance for AF3 to obtain a good MSA to build accurate structural models than the default MSA, which is particularly useful for hard monomer targets.

#### 3.1.2 Comparison of AF3 and AF2 on individual MSA-template combinations in tertiary structure prediction

A detailed comparison of AF3 and AF2 predictions using many MSA and template inputs across the CASP16 monomer targets is shown in Fig. 4a. Multiple different MSA and template inputs for each CASP16 monomer target were tested and the TM-scores of the top-ranked AF3 and AF2 models for each of them were compared. For most MSA-template combinations across almost all targets, the top-ranked AF3 models achieved higher TM-scores than their AF2 counter-parts, with average top-ranked TM-scores of 0.868 for AF3 and 0.852 for AF2. The difference is statistically significant according to one-sided Wilcoxon signed-rank test (p = 2.7987e-03 for TM-score), highlighting AF3’s improved ability to leverage evolutionary and template information in the same MSA-template input. Overall, the results indicate that AF3 provides a consistent and statistically significant gain over AF2 under the same MSA-template input.

**FIGURE 4.**
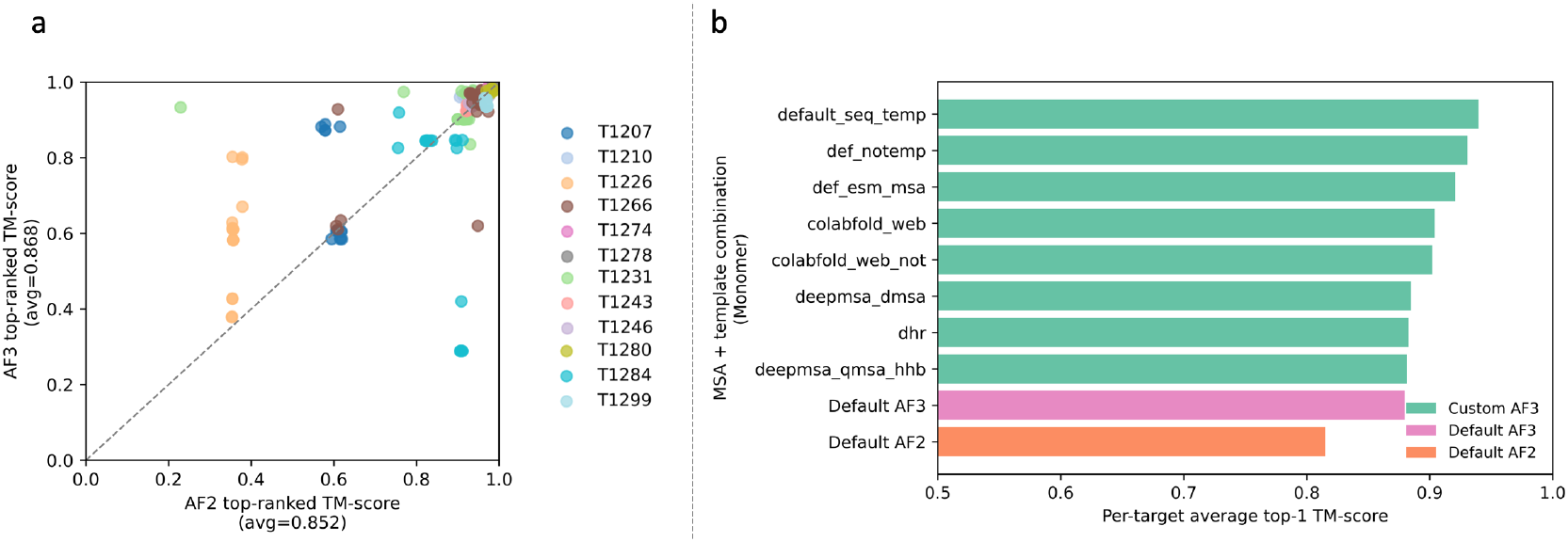
Head-to-head comparison of AF3 and AF2 as well as different MSA-template combination methods in monomer structure prediction. **a**. The comparison of TM-scores of top-ranked models from AF3 and AF2 using many MSA-template combinations as inputs across monomer targets. Each point denotes an input–target pair, where its x-coordinate corresponds to the score of the AF2 model and y-coordinate the score of the AF3 model. Colors indicating target identity. Multiple dots in the same color denote multiple MSA-template combinations for the same target. The dashed diagonal denotes equal performance. Mean TM-scores across all targets for all MSA-template combinations are shown in parentheses. **b**. Comparison of the average top-ranked TM-scores of different MSA-template generation methods for AF3 across 12 targets. Default AF3 MSA-template combination and default AF2 MSA-template combination are also included for comparison.

Moreover, as shown in Fig. 4a, in some cases, different MSA-template combinations for the same target (the dots of the same color) have a large variation of TM-scores (spread widely), while in other cases, they have a low variation (cluster closely), indicating that AlphaFold prediction for different targets has different levels of sensitivity to the difference in MSA and template input. In addition, AF3 and AF2 may have a different sensitivity to the same target. For instance, for hard target T1226, the scores of the AF3 models of different MSA-template combinations (orange dots) vary substantially from below 0.4 to 0.8, but those of AF2 models change little around a low score of 0.36.

Figure 4b compares different MSA–template generation methods using the average TM-score of their top-ranked models across the CASP16 monomer targets (see the description of each MSA-template generation method in Supplementary Table S1). Default AF3 outperforms Default AF2, with an average top-1 TM-score of 0.880 versus 0.815. Other MSA-template generation methods perform better than both default AF3 and default AF2. For instance, AF3 using the default AF2 MSA together with Custom templates retrieved from the PDB_sort90 database (denoted as default_seq_temp) achieves an average top-ranked TM-score of 0.940, exceeding the 0.880 of Default AF3 and the 0.815 of Default AF2. It is worth noting that, because the number of monomer targets (12) is relatively small, the difference between the MSA-template generation methods may not be statistically significant, the results show that AF3 benefits strongly from enriching its inputs with complementary MSAs and templates.

### 3.2 Quaternary Structure Prediction for Protein Multimers

#### 3.2.1 Comparison of Custom AF3, default AF3, and Custom AF2 on CASP16 multimer targets

We evaluate the performance of Custom AF3, default AF3, and Custom AF2 on 36 CASP16 multimer targets using six complimentary metrics, including TM-score for the global fold accuracy, DockQ, ICS, IPS, and QS_best scores for interface accuracy, and lDDT for local distance accuracy. The default AF3 used the standard MSA and template inputs provided by AlphaFold3 to generate between 425 and 5400 structural models per target, depending on the size of a target. The Custom AF3 generated the same total number of models, but used multiple Custom MSA–template combinations, producing 25–100 models for each combination. In both cases, models were ranked and selected for evaluation using the AF3 ranking score. Analogous to Custom AF3, Custom AF2 also employed all custom MSA–template combinations to generate a pool of structural models; to ensure a fair comparison, the same number of models (25–100) were randomly sampled for Custom AF2 from each MSA-template combination as Custom AF3, resulting in the same total number of structural models per target for Custom AF2. The models of Custom AF2 were ranked by the AF2 confidence score.

The average per-target performance in terms of multiple quality metrics shows that the Custom AF3 consistently outperforms default AF3 and Custom AF2, with particularly pronounced gains in DockQ score (0.550 for Custom AF3, 0.525 for default AF3, and 0.528 for Custom AF2) (Fig. 5a). Different from the CASP16 monomer targets, default AF3 does not outperform Custom AF2 on the CASP16 multimer targets. However, when the same diverse MSAs and templates are used, Custom AF3 performs consistently better than Custom AF2, indicating the importance of using complementary MSAs and templates to improve the accuracy of AlpahFold3. The average scores of the best models produced by each method are reported in Supplementary Table S5, which also demonstrate Custom AF3 performs best.

**FIGURE 5.**
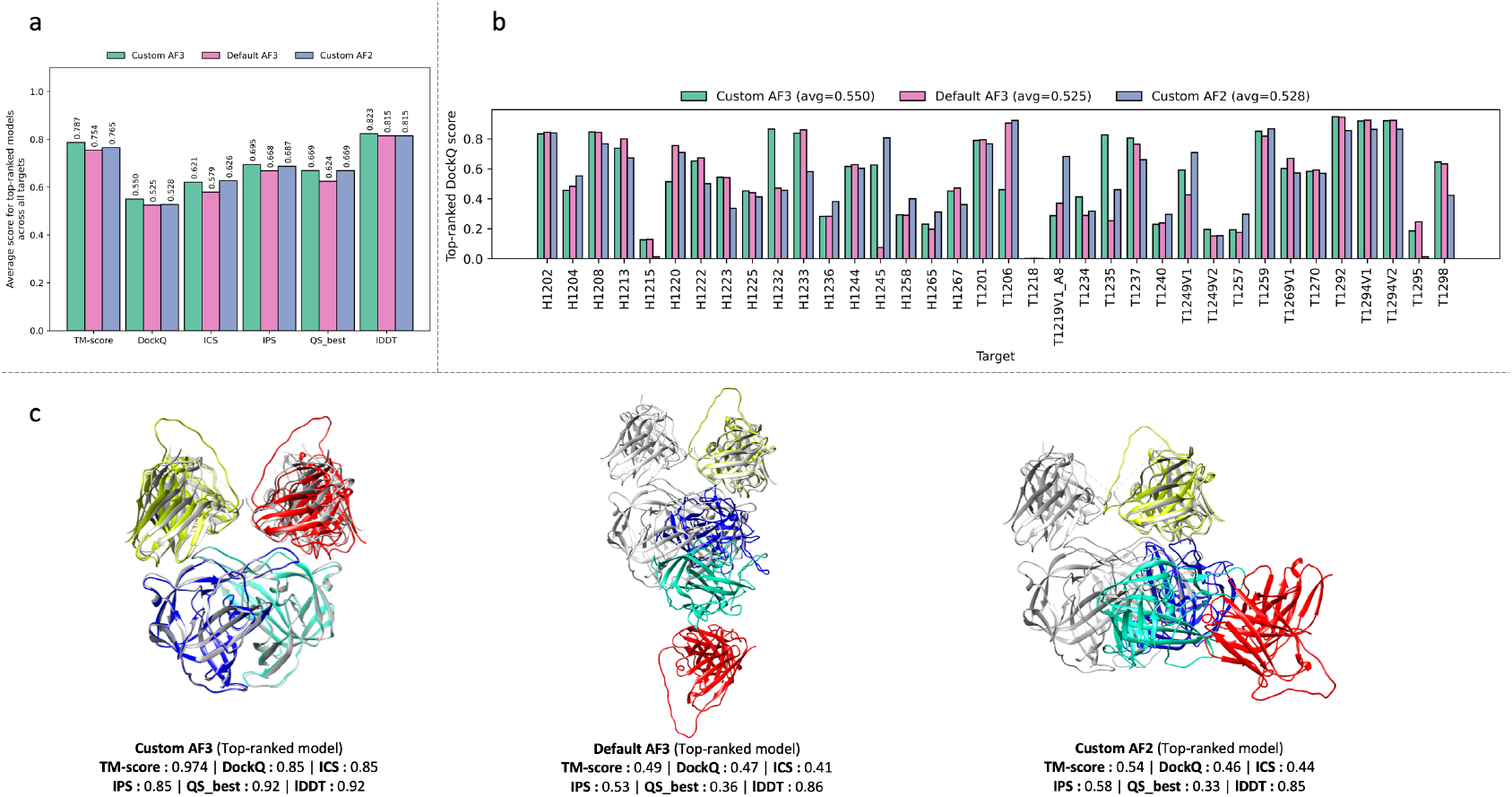
Comparative performance of Custom AF3, default AF3, and Custom AF2 on CASP16 multimer targets. **a**. Average per-target performance in terms of TM-score, DockQ, ICS, IPS, QS_best, and lDDT for the top-ranked model from each method. **b**. DockQ scores of top-ranked models of Custom AF3 (green), default AF3 (pink) and top-ranked Custom AF2 (blue) for 36 multimer targets. Mean DockQ scores across all targets are shown in parentheses. **c**. The top-ranked structural models for a heteromultimer target, H1232 (stoichiometry: A2B2), predicted by each method, in comparison with the native structure. The native structure is colored in silver, while the predicted structures are highlighted in rainbow colors. The six quality scores are indicated for each method.

Target-level comparisons of the top-ranked models show that Custom AF3 outperforms the other two methods on many targets, but there are also some targets where AF3 performs similarly to or worse than Custom AF2 or default AF3, highlighting substantial target-to-target variability (Fig. 5b).

Figure 5c shows a representative antibody target (H1232; stoichiometry A_2_B_2_) on which Custom AF3 substantially outperformed default AF3 and Custom AF2. Default AF3 generates a partially correct assembly (TM-score 0.49; DockQ 0.47; ICS 0.41; IPS 0.53; QS_best 0.36; lDDT 0.86), while Custom AF2 produces a nearly similar model with only a modest TM-score increase (+0.05). In contrast, Custom AF3 yields a highly accurate prediction (TM-score 0.974; DockQ 0.85; ICS 0.85; IPS 0.85; QS_best 0.92; lDDT 0.92), correctly recovering both the global quaternary structure and key interfacial interactions. Structurally, this A_2_B_2_ complex can be viewed as two A_1_B_1_ heterodimers arranged symmetrically. Across all three predictions, the internal A–B pairing within each A_1_B_1_ dimer is largely consistent (e.g., the yellow and blue components align well), indicating that heterodimer formation is not the primary challenge. Instead, the dominant difference lies in the relative placement of the two A_1_B_1_ dimers: Custom AF3 positions them in a symmetric, tightly packed configuration that completes the tetramer, whereas default AF3 and custom AF2 place the second A_1_B_1_ dimer with an incorrect translation/rotation, disrupting the inter-dimer contacts required for the A_2_B_2_ assembly and leading to substantially lower interface scores despite comparable local accuracy.

#### 3.2.2 Comparison of AF3 and AF2 on individual MSA-template combinations in quaternary structure prediction

Head-to-head comparisons between AF3 and AF2 using many MSA-template combinations across 36 CASP16 multimer targets show that AF3 achieves higher accuracy on most targets in both global quaternary arrangement and interface modeling (Fig. 6a). Specifically, AF3 attains a higher average top-ranked TM-score (0.765 of AF3 vs. 0.721 of AF2) and a higher average top-ranked DockQ score (0.515 of AF3 vs. 0.462 of AF2). These improvements are statistically significant by one-sided Wilcoxon signed-rank tests (p-value = 5.2272e-28 for TM-score; p-value = 2.6335e-51 for DockQ), indicating that AF3 more effectively exploits the same evolutionary and template inputs for multimer prediction.

**FIGURE 6.**
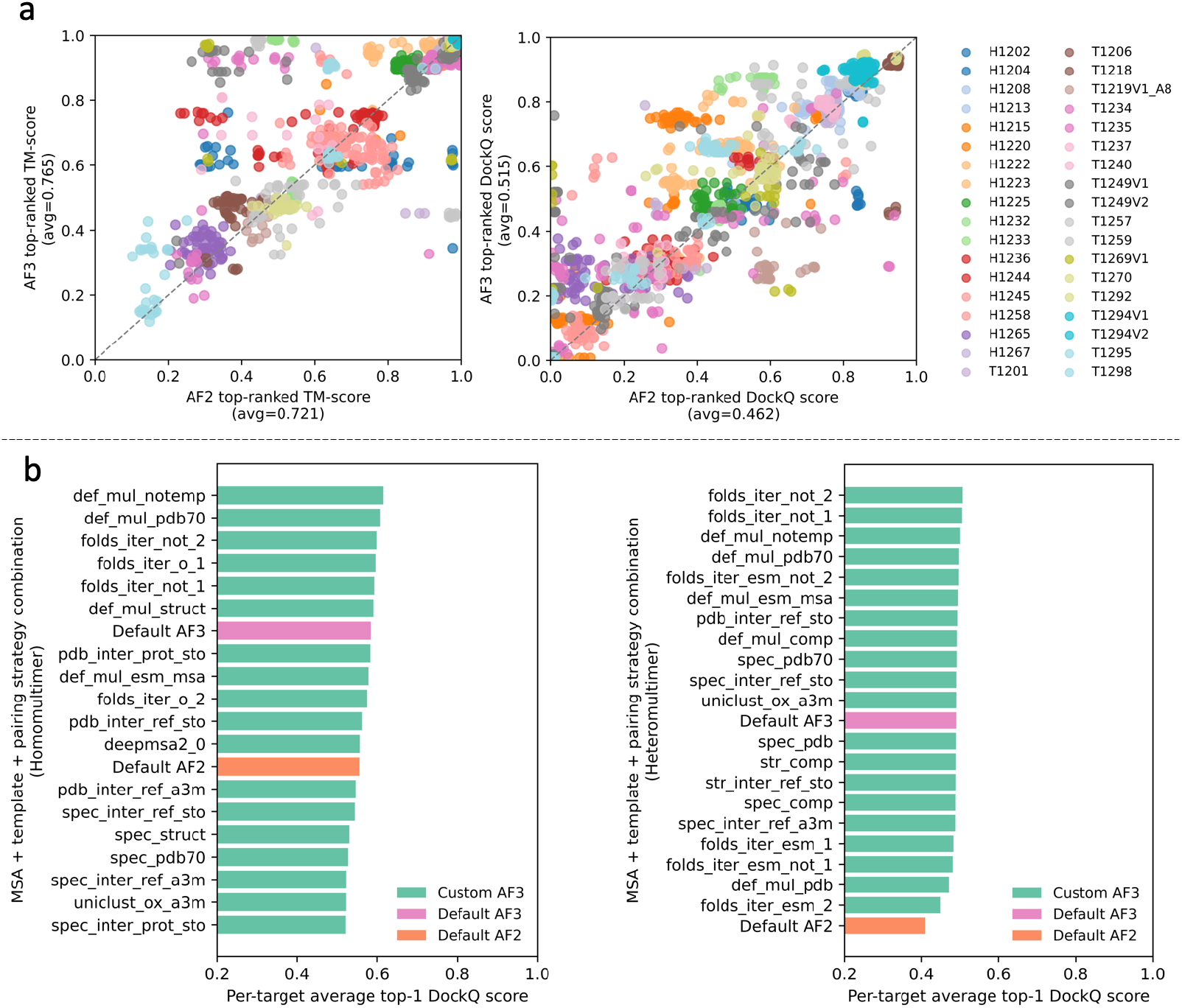
Head-to-head comparison of AF3 and AF2 as well as different MSA-template generation methods in multimer structure prediction. **a**. Comparison of TM-scores and DockQ scores of top-ranked models from AF3 and AF2 using multiple sets of MSA-template combinations as inputs across CASP16 multimer targets. Each point denotes an input–target pair, where its x-coordinate corresponds to the score of the AF2 model and y-coordinate the score of the AF3 model. Colors indicating target identity. Multiple dots in the same color denote multiple MSA-template combinations for the same target. The dashed diagonal denotes equal performance. Mean TM-scores and DockQ scores across all targets for all MSAs and template combinations are shown in parentheses. **b**. Comparison of performance of different MSA and template generation methods used with AF3, averaged over 15 homomultimeric targets (left) and 15 heteromultimeric targets (right) in terms of the DockQ score of top-ranked models. Default AF3 and default AF2 are also included as baseline for comparison.

Moreover, as shown in Fig. 6a, different MSA-template combinations for the same targets (denoted by the dots in the same color) sometime spread widely for AF3, AF2, or both, indicating they can cause substantial difference in the quality of generated models (TM-score or DockQ). Because it is still impossible to accurately assess the usability of a MSA-template input for quaternary structure prediction beforehand, it is important to explore multiple, complementary MSA-template combinations in model generation.

Figure 6b compares the performance of different MSA-template generation methods, summarized by the average per-target DockQ of top-1 models across 15 homomultimeric targets (left panel) and 15 heteromultimeric targets (right panel) (see the description of the MSA-template generation methods in Supplementary Tables S2 and S3). Six targets (four homomultimers: T1240, T1257, T1219V1_A8, and T1269V1; and two heteromultimers: H1232 and H1233) were excluded from this analysis because Foldseek-based MSA and corresponding template inputs were not generated during the 2024 CASP16 experiment. Across both panels, multiple MSA-template generation methods outperform default AF3, demonstrating that AF3 multimer prediction accuracy can be further improved by enriching MSA/template inputs.

For homomultimers, default AF3 achieves a higher average top-ranked DockQ score than default AF2 (0.584 vs. 0.555). Among the evaluated MSA–template generation methods, the best-performing method (`def_mul_notemp`) uses the default AlphaFold2 alignments without template information, which attains the highest average DockQ score of 0.616 when used with AF3. This is followed by `def_mul_pdb70`, which uses the default AlphaFold2 MSA with template information from PDB70. Notably, several of the next highest-ranking methods (e.g., `folds_iter_not_2`) are based on FoldSeek structure alignment-based MSAs.

For heteromultimers, DockQ scores are generally lower and more tightly clustered than (Fig. 6b, right), consistent with the greater difficulty of assembling non-identical subunits and their interfaces. Nevertheless, default AF3 substantially improves over default AF2 (0.490 vs. 0.410), and a Foldseek-based MSA generation method (i.e., `folds_iter_not_2`) achieve the best results. `folds_iter_not_2` MSA-template combination increases the average top-ranked DockQ score to 0.506.

The consistent presence of FoldSeek structure alignment-based MSA generation among the top-performing methods indicates that such MSAs provide reliable gains in AF3 prediction accuracy for both homomultimeric and hetero-multimeric complexes.

### 3.3 Structure Prediction for Protein–Ligand Complexes

We compared Custom AF3 and default AF3 on 286 CASP16 protein-ligand targets for which the native structures are available (Fig. 7). To evaluate protein–ligand complex predictions, we used ligand RMSD to assess global binding pose accuracy, lDDT_PLI to quantify local accuracy at the protein–ligand interface, backbone RMSD (BB-RMSD) to measure receptor conformational changes, and lDDT-LP to evaluate local structural quality in the ligand-proximal region. The default AF3 used the standard MSA and template inputs provided by AF3 to generate between 1100 and 3200 structural models per target. Custom AF3 generated the same total number of models, but relied on custom multiple MSA–template combinations, producing 100 models for each combination. In both cases, models were ranked and selected for evaluation using the AF3 ranking score.

**FIGURE 7.**
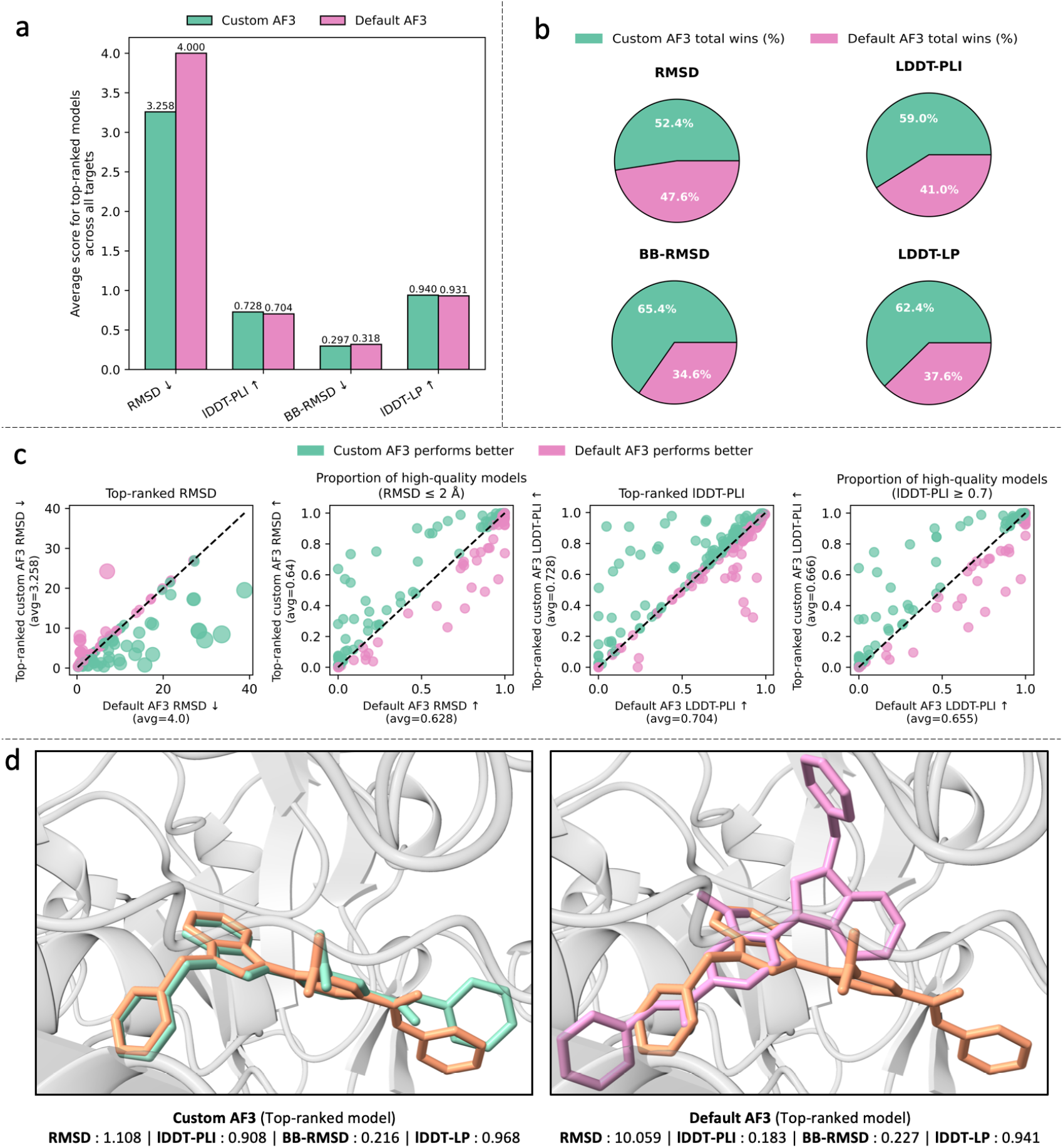
Comparison of Custom AF3 and default AF3 across CASP16 protein-ligand targets. **a**. Average performance across targets in terms of ligand RMSD, lDDT-PLI, backbone RMSD (BB-RMSD) and lDDT-LP. **b**. Head-to-head comparison summarizing the fraction of targets where Custom AF3 outperforms or underperforms default AF3 according to each of the 4 evaluation metrics. **c**. Scatter bubble plots comparing, from left to right, top-ranked ligand RMSD, the proportion of high-quality models based on ligand RMSD (RMSD ≤ 2 Å), top-ranked lDDT-PLI, and the proportion of high-quality models based on lDDT-PLI (lDDT-PLI ≥ 0.7) between Custom AF3 (green) and default AF3 (pink). Each point represents a target, where x-coordinate corresponds to the score of the top-ranked AF2 model and y-coordinate the score of the top-ranked AF3 model; dashed lines indicate equal performance. **d**. A representative example (target L3157) illustrating ligand placement. The experimental ligand (orange) is overlaid with predictions from Custom AF3 (green, left) and default AF3 (pink, right).

The average structural accuracy of top-ranked models of Custom AF3 and default AF3 across all targets in terms of the four metrics is reported in Fig. 7a. Custom AF3 consistently improves global and local accuracy relative to default AF3. In particular, Custom AF3 yields a substantially lower average RMSD (3.258 Å of Custom AF3 vs 4 Å of default AF3) and BB-RMSD (0.297 vs 0.318), indicating more accurate global folds and backbone conformations achieved by using multiple MSA-template combinations. At the same time, it achieves higher lDDT-PLI (0.728 vs 0.704) and lDDT-LP (0.94 vs 0.931) scores, reflecting improved local structural fidelity at protein–ligand interface and ligand-proximal regions. The results show that using diverse MSA and templates of a receptor protein with AF3 can improve the prediction of its interaction with ligands, even though the ligand input information is the same.

To assess the consistency of these improvements across individual targets, we quantified the proportion of cases in which each method outperforms the other (Fig. 7b). Custom AF3 has a higher fraction of winning cases in terms of all evaluated metrics, including RMSD, lDDT-PLI, BB-RMSD, and lDDT-LP. The advantage is particularly pronounced for backbone accuracy and ligand-proximal local quality, indicating that the gains observed in average performance are not driven by a small number of outliers but instead reflect robust improvements across the benchmark.

A target-wise comparison between Custom AF3 and default AF3 is plotted in Fig. 7c, including the comparison of RMSD of top-ranked models, proportion of high-quality models in terms of RMSD, lDDT-PLI of top ranked models, and proportion of high-quality models in terms of lDDT-PLI. For top-ranked RMSD, the majority of targets lie below the diagonal, indicating consistently lower errors for Custom AF3. Conversely, for top-ranked lDDT-PLI, most targets lie above the diagonal, reflecting improved accuracy at the protein–ligand interface.

In addition, Custom AF3 increases the overall prevalence (proportion) of high-quality models within the generated ensembles. Specifically, the average fraction of models achieving RMSD ≤ 2 Å increases from 0.628 to 0.64, while the average fraction achieving lDDT-PLI ≥ 0.7 increases from 0.655 to 0.666.

Fig. 7d illustrates a representative example illustrates the practical impact of these improvements. For the target L3157, Custom AF3 produced a highly accurate ligand pose with substantially better scores across all 4 metrics (RMSD : 1.108 Å, lDDT-PLI : 0.908, BB-RMSD : 0.216, lDDT-LP : 0.968), whereas default AF3 mispositioned the ligand, resulting in poorer scores, particularly for RMSD and lDDT-PLI (RMSD : 10.059 Å, lDDT-PLI : 0.183, BB-RMSD : 0.227, lDDT-LP : 0.941).

## CONCLUSION

In this study, we show that systematic MSA and template engineering yields consistent and substantial improvements in AlphaFold3 predictions across monomeric proteins, multimeric complexes, and protein–ligand assemblies. By integrating diverse evolutionary signals through complementary search strategies, our framework improves backbone accuracy, interface modeling, and ligand placement, without altering the underlying model architecture of AlphaFold3. These findings emphasize that, even as structure prediction evolves toward unified foundation models, the quality and diversity of evolutionary context remain central determinants of predictive performance. The tools and guidelines presented here establish a practical and broadly applicable framework for strengthening biomolecular structure modeling, helping to further advance next-generation structural biology workflows.

## 4 ACKNOWLEDGEMENTS

We thank CASP16 organizers and assessors for making the CASP16 data available. This work is partially supported by two NIH grants [R01GM093123 and R01GM146340], NSF grants [DBI2308699, IOS2525780], and a Department of Energy grant [DE-SC0026121].

## 5 AUTHOR CONTRIBUTIONS

J.C. conceived the project. J.C., P.N., J.L. designed the experiment. P.N. and J.L. performed the experiment and collected the data. P.N., J.L. and J.C. analyzed the data. P.N., J.L. and J.C. wrote the manuscript.

## 7 DATA AVAILABILITY

The protein structures of the CASP16 monomer/complex/protein-ligand targets are available at https://predictioncenter.org/download_area/CASP16/targets/. All MSAs and templates used in this study, as well as the structural models generated for evaluation are publicly available via Harvard Dataverse at https://doi.org/10.7910/DVN/KGWHRE.

## 8 CODE AVAILABILITY

The code of Custom AF3 is publicly available at: https://github.com/BioinfoMachineLearning/CAF3.

