## Supplementary data for "Improving AlphaFold3 by Engineering MSA and Template Inputs"

April 22, 2026

Table S1: Summary of MSA and template generation methods for monomers.

| Name of MSA | MSA generation tool | Sequence database | Template database |
|---|---|---|---|
| default_seq_temp | HHblits, JackHMMER | UniRef90, UniRef30 & BFD, MGnify clusters | PDB_sort90 |
| def_notemp | HHblits, JackHMMER | UniRef90, UniRef30 & BFD, MGnify clusters | No template |
| def_esm_msa | HHblits, JackHMMER, ESM2 | UniRef90, UniRef30 & BFD, MGnify clusters | pdb70 |
| colabfold_web | MMseq2 | ColabFold DB | pdb70 |
| colabfold_web_not | MMseq2 | ColabFold DB | No template |
| deepmsa_dmsa | DeepMSA2 | TaraDB, Metaclust, MetaSourceDB, and JGIclust | pdb70 |
| dhr | DHR | UniRef90 | pdb70 |
| deepmsa_qmsa_hhb | DeepMSA2 | TaraDB, Metaclust, MetaSourceDB and JGIclust | pdb70 |

Table S2: Summary of MSA and template generation methods for homo-multimers..

| **Name** | $\mathbf{MSA_{paired}}$ | **Interaction Source** | **Template** |
|---|---|---|---|
| def_mul_notemp | Default | - | No template |
| def_mul_pdb70 | Default | - | pdb70 |
| folds_iter_not_2 | Structural alignments from pdb_complex & AlphaFoldDB + subunit MSA | Identifier from alignments | No template |
| folds_iter_o_1 | Structural alignments from pdb_complex & AlphaFoldDB + subunit MSA | Identifier from alignments | Structural template |
| folds_iter_not_1 | Structural alignments from pdb_complex & AlphaFoldDB + subunit MSA | Identifier from alignments | No template |
| def_mul_struct | Default | - | pdb_complex |
| def_mul_esm_msa | Default + ESM-MSA | - | pdb_seqres |
| pdb_inter_prot_sto | Uniprot | PDB Code | pdb_seqres |
| folds_iter_o_2 | Structural alignments from pdb_complex & AlphaFoldDB + subunit MSA | Identifier from alignments | Structural template |
| pdb_inter_ref_sto | UniRef90 | PDB code | pdb_seqres |
| deepmsa2_0 | Top-ranked MSA from DeepMSA2 | DeepMSA2 | pdb_seqres |
| pdb_inter_ref_a3m | UniRef30 | PDB Code | pdb_seqres |
| spec_inter_ref_sto | UniRef90 | Species annotation | pdb_seqres |
| spec_struct | UniRef30 | Species annotation | pdb_complex |
| spec_pdb70 | UniRef30 | Species annotation | pdb70 |
| spec_inter_ref_a3m | UniRef30 | Species annotation | pdb_seqres |
| uniclust_ox_a3m | UniClust30 | Species annotation | pdb_seqres |
| spec_inter_prot_sto | UniProt | Species annotation | pdb_seqres |

Table S3: Summary of MSA and template generation methods for hetero-multimers.

| **Name** | **MSA$_{\mathbf{paired}}$** | **Interaction Source** | **MSA$_{\mathbf{unpaired}}$** | **Template** |
|---|---|---|---|---|
| folds_iter_not_2 | Structural alignments from pdb_complex & AlphaFoldDB | Identifier from alignments | Subunit MSA + structural alignments | No template |
| folds_iter_not_1 | Structural alignments from pdb_complex & AlphaFoldDB | Identifier from alignments | Subunit MSA + structural alignments | No template |
| def_mul_notemp | UniProt | Species annotation | Subunit MSA | No template |
| def_mul_pdb70 | UniProt | Species annotation | Subunit MSA | pdb70 |
| folds_iter_esm_not_2 | Structural alignments from ESM-Atlas | PDB code | Subunit MSA + structural alignments | No template |
| def_mul_esm_msa | UniProt | Species annotation | Subunit MSA + ESM-MSA | pdb_seqres |
| pdb_inter_ref_sto | UniRef90 | PDB code | Subunit MSA | pdb_seqres |
| def_mul_comp | UniProt | Species annotation | Subunit MSA | pdb_complex |
| spec_pdb70 | UniRef30 | Species annotation | Subunit MSA | pdb70 |
| spec_inter_ref_sto | UniRef90 | Species annotation | Subunit MSA | pdb_seqres |
| uniclust_ox_a3m | UniClust | Species annotation | Subunit MSA | pdb_seqres |
| spec_pdb | UniRef30 | Species annotation | Subunit MSA | PDB_sort90 |
| str_comp | UniRef90 | STRING | Subunit MSA | pdb_complex |
| spec_comp | UniRef30 | Species annotation | Subunit MSA | pdb_complex |
| spec_inter_ref_a3m | UniRef90 | STRING | Subunit MSA | pdb_seqres |
| folds_iter_esm_1 | Structural alignments from ESM-Atlas | Identifier from alignments | Subunit MSA + structural alignments | Structural template from ESM-Atlas |
| folds_iter_esm_not_1 | Structural alignments from ESM-Atlas | Identifier from alignments | Subunit MSA + structural alignments | No template |
| def_mul_pdb | UniProt | Species annotation | Subunit MSA | PDB_sort90 |
| folds_iter_esm_2 | Structural alignments from ESM-Atlas | Identifier from alignments | Subunit MSA + structural alignments | Structural template from ESM-Atlas |

Table S4: Average performance across 12 CASP16 monomeric targets. For each metric and statistic, the best-performing method is highlighted in bold.

| Metric | Method<br>Statistic | Custom AF3 | Default AF3 | Custom AF2 |
|---|---|---|---|---|
| TM-score | Best ( ↑ ) | **0.959** | 0.896 | 0.948 |
| | Top-ranked ( ↑ ) | **0.937** | 0.882 | 0.869 |
| GDT-TS | Best ( ↑ ) | **0.929** | 0.864 | 0.911 |
| | Top-ranked ( ↑ ) | **0.895** | 0.839 | 0.824 |
| GDC-SC | Best ( ↑ ) | **0.641** | 0.597 | 0.595 |
| | Top-ranked ( ↑ ) | **0.579** | 0.561 | 0.519 |
| lDDT | Best ( ↑ ) | **0.866** | 0.816 | 0.839 |
| | Top-ranked ( ↑ ) | **0.838** | 0.798 | 0.775 |

Table S5: Average per-target performance across 36 CASP16 multimeric targets. For each metric and statistic, the best-performing method is highlighted in bold.

| Metric | Method<br>Statistic | Custom AF3 | Default AF3 | Custom AF2 |
|---|---|---|---|---|
| TM-score | Best ( ↑ ) | **0.887** | 0.875 | 0.880 |
| | Top-ranked ( ↑ ) | **0.787** | 0.754 | 0.765 |
| DockQ | Best ( ↑ ) | **0.711** | 0.666 | 0.673 |
| | Top-ranked ( ↑ ) | **0.550** | 0.525 | 0.528 |
| ICS | Best ( ↑ ) | **0.775** | 0.746 | 0.746 |
| | Top-ranked ( ↑ ) | 0.621 | 0.579 | **0.626** |
| IPS | Best ( ↑ ) | **0.817** | 0.803 | 0.798 |
| | Top-ranked ( ↑ ) | **0.695** | 0.668 | 0.687 |
| QS | Best ( ↑ ) | **0.840** | 0.810 | 0.822 |
| | Top-ranked ( ↑ ) | **0.669** | 0.624 | 0.669 |
| lDDT | Best ( ↑ ) | **0.855** | 0.849 | 0.846 |
| | Top-ranked ( ↑ ) | **0.823** | 0.815 | 0.815 |
